# A Rarefaction-Based Extension of the LDM for Testing Presence-Absence Associations in the Microbiome

**DOI:** 10.1101/2020.05.26.117879

**Authors:** Yi-Juan Hu, Andrea Lane, Glen A. Satten

## Abstract

**Background:** Many methods for testing association between the microbiome and covariates of interest (e.g., clinical outcomes, environmental factors) assume that these associations are driven by changes in the relative abundance of taxa. However, these associations may also result from changes in which taxa are present and which are absent. Analyses of such presence-absence associations face a unique challenge: confounding by library size (total sample read count), which occurs when library size is associated with covariates in the analysis. It is known that *rarefaction* (subsampling to a common library size) controls this bias, but at the potential cost of information loss as well as the introduction of a stochastic component into the analysis. Currently, there is a need for robust and efficient methods for testing presence-absence associations in the presence of such confounding, both at the community level and at the individual-taxon level, that avoid the drawbacks of rarefaction.

**Methods:** We have previously developed the linear decomposition model (LDM) that unifies the community-level and taxon-level tests into one framework. Here we present an extension of the LDM for testing presence-absence associations. The extended LDM is a non-stochastic approach that repeatedly applies the LDM to *all* rarefied taxa count tables, averages the residual sum-of-squares (RSS) terms over the rarefaction replicates, and then forms an *F*-statistic based on these average RSS terms. We show that this approach compares favorably to averaging the *F*-statistic from *R* rarefaction replicates, which can only be calculated stochastically. The flexible nature of the LDM allows discrete or continuous traits or interactions to be tested while allowing confounding covariates to be adjusted for.

**Results:** Our simulations indicate that our proposed method is robust to any systematic differences in library size and has better power than alternative approaches. We illustrate our method using an analysis of data on inflammatory bowel disease (IBD) in which case samples have systematically smaller library sizes than controls.

**Conclusions:** The rarefaction-based extension of the LDM performs well for testing presenceabsence associations and should be adopted even when there is no obvious systematic variation in library size.

## Background

Ecologists often view metagenomic count data in one of two ways when testing for association between the microbiome and covariates of interest such as clinical outcomes or environmental factors. One is to treat the counts as quantitative (i.e., analyze as relative abundance data); the other is to discretize the count data to indicate only whether a taxon is present or absent in a sample. Although the first approach is probably more common in the medical literature, it is also possible that associations are driven by changes in which taxa are present in a sample. For example, in the human gut, increased species richness are known to be associated with more stable ecosystems [1, 2], which tend to be resistant to environmental pressures such as diet, antibiotic use and pathogen invasion [3, 4]. In contrast, a healthy vaginal microbiome is often characterized by low diversity that is dominated by *lactobacilli* [5]. Analyses based on relative abundances are probably more reasonable when common taxa are expected to play a dominant role, while analyses based on presence or absence of taxa may perform better when rare taxa are most important. Presence-absence analyses may also be more robust to extraction and amplification biases that occur in all microbiome sequencing experiments [6] and that plague most analyses of relative abundance data.

The most commonly used method for presence-absence analyses is PERMANOVA using a presence-absence dissimilarity measure. The unweighted UniFrac [7, 8] and Jaccard distances are most frequently adopted, and rarefaction to a common library size is often applied to preemptively control for any differences in library size. As with all PERMANOVA analyses, this approach only gives community-level (sometimes referred to as *global*) associations, not associations with individual taxa.

The LDM [9] is based on linear models that treat (possibly transformed) relative abundances of taxa as the outcome. A simple extension of the LDM for testing presence-absence associations would be to replace the relative abundance data with presence-absence data (and suppress any further transformation of the data). The resulting tests will thus inherit the useful features of the original LDM, namely providing unified testing of global and taxon-level associations, accommodating both continuous and discrete covariates as well as interaction terms to be tested either singly or in combination, allowing for adjustment of confounding covariates, and using permutation-based *p*-values that can control for between-sample correlation (e.g., matched-set data [10]). Since the LDM permutes covariates rather than the outcome (or residuals of the outcome), the binary nature of the outcome poses no special difficulties. Many recently proposed methods for testing association of the microbiome with covariates of interest explicitly assume that the association is driven by changes in relative abundance of taxa, and so cannot be applied to presence-absence data; these methods include metagenomeSeq [11], ANCOM [12, 13], and ALDEX2 [14].

All analyses of presence-absence data face the challenge of confounding by library size (i.e., sequencing depth), if library size varies with covariates of interest (e.g., disease status). Even though modern 16S rRNA gene sequencing typically yields large library sizes, under-sampling of rare taxa may still occur. Without additional information, it is impossible to distinguish between a taxon that is completely absent from the biological specimen, and a taxon that is actually present but has no reads in a particular measurement because of under-sampling. For this reason, if library size is associated with any covariate, that covariate may show a spurious association with presence or absence of rare taxa. In theory, confounding by library size may affect the analysis of relative abundance data as well, but the impact is typically much smaller.

Ecologists frequently use rarefaction [15, 16] to overcome the potential confounding effects of library size. Rarefaction corresponds to subsampling reads to a common *rarefaction depth*, which is often the lowest observed library size (after removing outliers). As rarefaction produces data with a single library size, there will be no covariates that are associated with library size. On the other hand, rarefaction typically results in a substantial loss of reads and hence statistical power. Multiple rarefaction [16] has been proposed to recover some of the information lost through rarefaction. However, it remains unclear how to aggregate the information from multiple rarefied datasets, and how many rarefactions recover sufficient information.

Simple taxon-level presence-absence analyses using rarefied data can also be conducted using standard statistical methods, as long as these methods can correctly handle sparse counts. For example, Fisher’s exact test can be used to test the association between a categorical trait and presence or absence of a single taxon. This is roughly analogous to testing taxon-level association using the Wilcoxon (or Kruskal-Wallis) test for relative abundance data. For more complex situations (e.g., with confounding covariates), exact logistic regression could be used, although this can be computationally expensive. Further, there remains the question of how to construct a community-level test from the results at each taxon. For these reasons, and because real microbiome experiments nearly always have potentially confounding covariates, the only methods for “binary” data analysis we consider here are based on the LDM.

We propose here an extension of the LDM for analyzing presence-absence data that accounts for the variation of library size. In the methods section, we will adopt rarefaction as our choice of normalization. We first consider two ways to combine multiple rarefactions, and then select the approach that allows us to combine information over all possible rarefied taxa count tables. In the results section, we present the simulation studies and the application to a real study on inflammatory bowel diseases (IBD). We conclude with a discussion section.

## Methods

The LDM is a linear model in which the covariates (metadata) are summarized in a design matrix *X* (where the rows correspond to *n* samples and the columns correspond to the covariates). We may partition *X* by columns into *K* groups (which we call “submodels”) such that *X* = (*X*_1_, *X*_2_, …, *X*_*K*_), where each column vector or *n* × *J*_*k*_ matrix *X*_*k*_ denotes a variable or set of variables we wish to test jointly. For example, *X*_*k*_ may consist of indicator variables for levels of a single categorical variable, or a group of potential confounders that we wish to adjust for simultaneously (such as a set of variables describing smoking history). The LDM makes the columns of *X orthonormal*.

We now consider how to combine information on rarefaction replicates using the LDM. Let *Y* be the (original) taxa count table of read counts and *Y* ^(*r*)^ the count table after the *r*th rarefaction. We convert *Y* ^(*r*)^ into a presence-absence (i.e., binary) matrix *B*^(*r*)^ = 𝕀(*Y* ^(*r*)^ *>* 0), where 𝕀(.) is the indicator function that performs element-wise discretization of its matrix argument. The LDM uses an *F*-statistic to test both taxon-specific and global effects. The *F* statistic (we drop the “minus one” term as well as a multiplier consisting of degrees of freedom in the original *F* statistic, since the significance is assessed via permutation) for submodel *k* is the ratio of residual sum-of-squares (RSS) terms for the model that excludes submodel *k* (in the numerator) or includes submodel *k* (in the denominator). One way to combine information over rarefaction replicates would be to use the average of the *F* statistic over the replicates as a test statistic. We also consider a second possible way to combine information, by separately averaging the RSS terms in the numerator and denominator, and then use the ratio of these averages as a test statistic. Note that in either case, we obtain both taxon-level and global tests, by permuting covariates as in the original LDM.

A priori, it would seem that averaging the *F* statistics is the safer route, since the average is taken after the final step of the analysis of a single rarefaction replicate. However, two factors argue for averaging the individual RSS terms. First, if we were considering the same question but using PERMANOVA, averaging the individual RSS terms corresponds to averaging the (element-wise squared) distance matrix over rarefaction replicates and then using the averaged matrix to calculate the PERMANOVA *F* statistic. This strategy is implemented, for example, in the R package `vegan` where the `avgdist` function calculates the average distance matrix over rarefaction replicates. Second and perhaps more importantly, for the LDM, the averages of the RSS terms over all rarefaction replicates can be easily calculated in closed forms. As a result, it is not necessary to perform any actual rarefactions to implement the second way for the LDM.

To see how the averages of the RSS terms in the LDM over all rarefaction replicates can be calculated in closed forms, we note that the *F* statistic in the LDM for the effect of taxon *j* on submodel *k* has the form *F*_*kj*_ = RSS_*kj*1_*/*RSS_*kj*2_, where RSS_*kj*1_ and RSS_*kj*2_ are the numerator and denominator RSS terms, respectively. Each RSS has the form

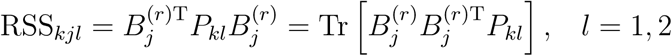

where 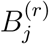 is the *j*th column of the matrix *B*^(*r*)^ and *P*_*kl*_ (*l* = 1, 2) are the projection matrices whose formulas can be found in [9].

The sampling distribution for the read count of taxon *j* in sample *i* sampled in rarefaction 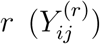 follows the hypergeometric distribution given the original library size of sample *i* (*N*_*i*_), the rarefaction depth (*N*_0_), and the original read count of taxon *j* in sample *i* (*Y*_*ij*_), as illustrated in Table 1. Thus, 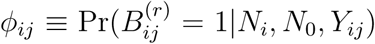 is one minus the probability that 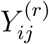 receives zero count:

**Table 1.** The 2 × 2 table illustrating the hypergeometric sampling

| In rarefaction $r$ | taxon $j$ | All other taxa | |
| --- | --- | --- | --- |
| Sampled for sample $i$ | $Y_{ij}^{(r)}$ | $N_0 - Y_{ij}^{(r)}$ | $N_0$ |
| Not sampled | $Y_{ij} - Y_{ij}^{(r)}$ | $N_i - N_0 - Y_{ij} + Y_{ij}^{(r)}$ | $N_i - N_0$ |
| | $Y_{ij}$ | $N_i - Y_{ij}$ | $N_i$ |

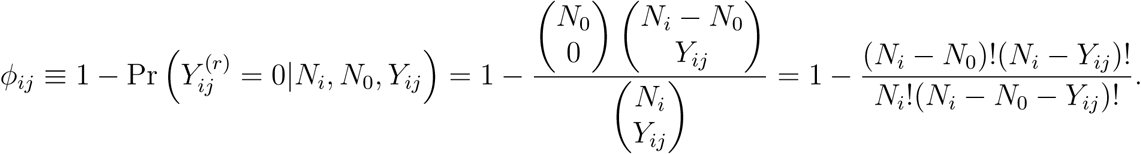

Since rarefactions are performed independently for each sample, it is not hard to calculate the expected value of RSS_*kjl*_ under repeated rarefactions, which is

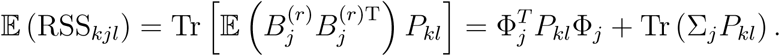

where Φ_*j*_ is the column vector with *i*th element *ϕ*_*ij*_ and ∑_*j*_ is a diagonal matrix with (*i, i*)th element given by *ϕ*_*ij*_(1 *–ϕ*_*ij*_). The expected value of RSS_*kjl*_ gives exactly the average of RSS_*kjl*_ over all rarefaction replicates. Note that it is still necessary to specify a rarefaction depth *N*_0_ even though the average can be computed without actually performing any rarefactions.

## Results

### Simulation studies

We created simulated data for 50 cases and 50 controls in which a binary confounder and the case-control status were associated with presence or absence of taxa in the microbiome. We based our simulations on the relative abundances of the 856 taxa reported in a study of the microbiome of the upper respiratory tract (URT) [17]. To generate taxa that were differentially present in cases but not controls, we uniformly selected 100 taxa (after excluding taxa with relative abundance *>* 1%) to be potentially associated with the case-control status *D*. Using the same approach, we independently selected a second set of 100 taxa to be associated with the confounder *C*, which was a binary variable with 70% “success” rate in controls but only 30% in cases; note that the sets of taxa associated with *D* and *C* may overlap. Denote the mean relative abundances of all taxa estimated from the URT data by the vector *π*, the entries of which are all positive, and denote the relative abundances of all taxa in sample *i* by the vector *π*_*i*_. For each sample, we initially set *π*_*i*_ = *π*; if the sample is a case, we then set the entries for each taxon selected to be associated with *D* to 0 with probability *β* (*∈* [0, 1]). This operation was done independently for each taxon. We used the value of *β* as a measure of the *effect size*, and noted that *β* = 0 corresponds to the null hypothesis of no association. For a sample with *C* = 1, we further set the entries in *π*_*i*_ to 0, independently for each taxon selected to be associated with *C*, with fixed probability 0.5. To ensure that all entries in *π*_*i*_ sum up to 1, for each sample we increased the probability mass of the most abundant taxon by the total mass that had been set to 0; since the most abundant taxon has sufficient probability mass to ensure it is always present, this operation does not change its presence-absence status.

Given *π*_*i*_, we then sampled read count data using the Dirichlet-Multinomial (DM) model. We set the overdispersion parameter to 0.02, which was also estimated from the URT data. We considered two scenarios for library size: a low-throughput setting with a mean of 1.5K reads as observed in the URT data, and a high-throughput setting with mean read count 10K. We used these values for controls and varied them for cases so as to introduce systematic differences in library size. Given a mean library size *µ*, the library size for each sample was drawn from *N* (*µ, µ/*3). Unless otherwise stated, the sampled library size was truncated at 2,500 in the high-throughput setting and 500 in the low-throughput setting. The truncation values were used as the rarefaction depth; in this way, no samples were discarded due to rarefaction (assuming that samples with problematic library sizes have been removed a priori). To analyze these data using the LDM, we took the design matrix *X* = (*C, D*), where we entered the confounder in the first column of *X* so that its effect was controlled when we estimated the effect of *D*.

We applied the proposed method that extends the LDM by averaging the RSS terms over all rarefactions, referred to as LDM-A. As a benchmark, we also applied the LDM that averages the *F* statistic over *R* rarefactions, referred to as LDM-F(*R*). In some studies, we included the LDM applied to unrarefied presence-absence matrix *B* = 𝕀(*Y >* 0), referred to as LDM-UR. We also evaluated the strategy that simply adjusted the library size as a covariate in the LDM applied to unrarefied data, referred to as LDM-L. We evaluated the size (i.e., type I error) and power for testing the global hypothesis of no case-control differences after adjusting for the confounder *C*; the nominal significance level was set to 0.05. We also assessed empirical sensitivity and empirical FDR for detecting individual taxa that were differentially present between cases and controls after adjusting for *C*; the nominal FDR was set to 10%. Results for size were based on 10,000 simulation replicates; all other results were based on 1,000 replicates.

#### Comparing different versions of the LDM in the presence of systematic differences in library size

We compared the performance of LDM-A, LDM-F(*R*) where *R* = 1, 5, LDM-UR, and LDM-L; the results of size are displayed in Figure 1. The performance of LDM-UR worsens as the difference in library size between cases and controls increases, while the other methods all control size in these simulations. It should be noted that LDM-L controls size in this setting probably because the library size is essentially a simple binary covariate (i.e., differing in mean in cases and controls); presumably LDM-L would not work well in more complex settings if the functional form of the dependence on library size were misspecified. To demonstrate that we have induced a substantial confounding effect, we show in Figure S1 that the size of LDM-A is as large as 0.2 when the confounder is not controlled for.

**Figure 1.**
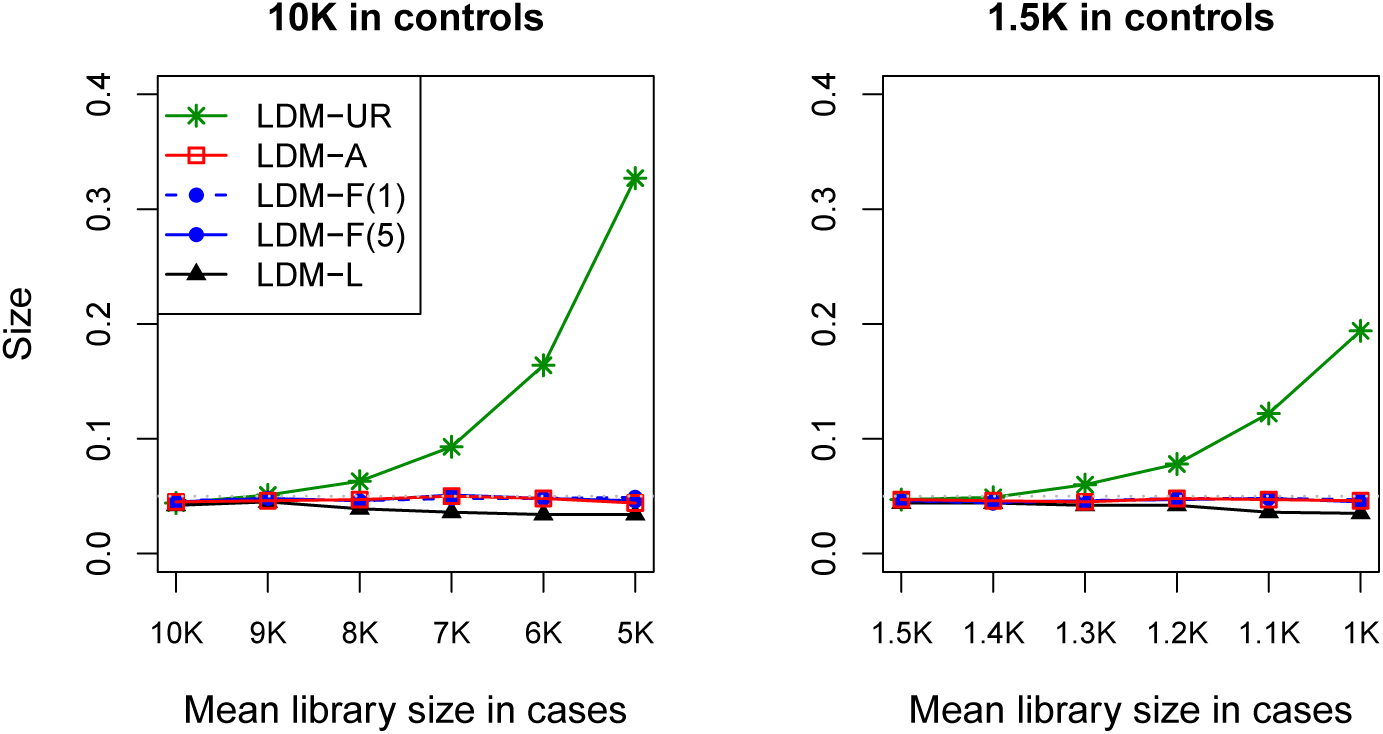
Size of the global test, when there are differential library sizes in cases and controls. The gray dotted line represents the nominal significance level of 0.05.

Figure 2 shows the power of LDM-A, LDM-F(5), and LDM-L for two settings with differential library sizes: (1) 10K and 5K for the mean library sizes of controls and cases, respectively, and (2) 1.5K and 1K. LDM-A yields the highest power (Figure 2, upper panel) for testing the global hypothesis and the highest sensitivity (Figure 2, middle panel) for detecting differentially present taxa, when compared to LDM-F(5) and LDM-L; all methods control FDR (Figure 2, lower panel). While LDM-F(5) only loses a little power and sensitivity to LDM-A, LDM-L loses substantial power and sensitivity. Note that we used 5 rarefactions for LDM-F(*R*) because we found that the power of LDM-F(*R*) nearly stabilizes after 5 rarefactions in our simulation studies (Figure S2). However, this pattern of power over rarefactions depends the underlying data; our results only suggest that at least 5 rarefactions would be needed for the LDM-F(*R*) method in general. Power results for LDM-UR are not shown as this method does not control size.

**Figure 2.**
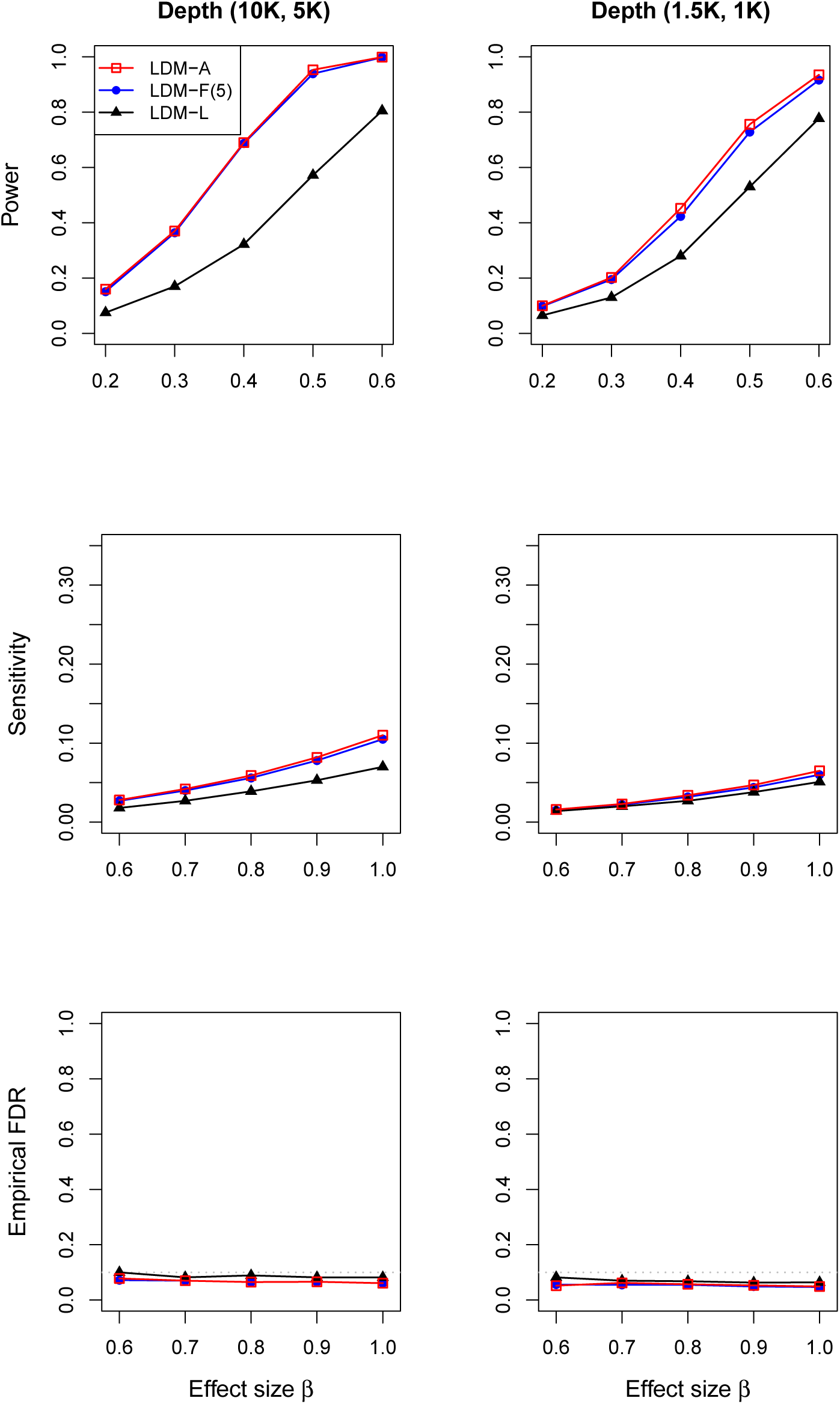
Power of the global test, and sensitivity and empirical FDR of the taxon-level tests.

**Figure 3.**
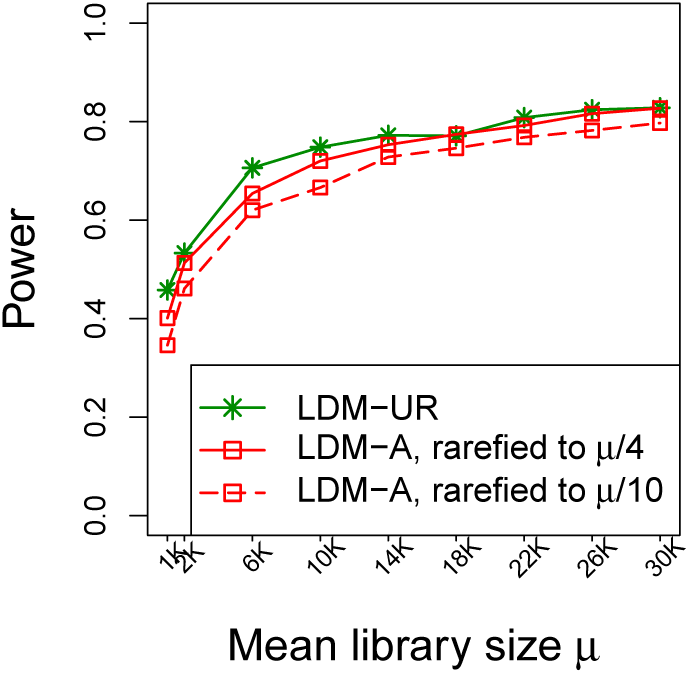
Power of the global test with and without rarefaction, when there are no systematic differences in library size between cases and controls. The effect size *β* was fixed at 0.4.

#### The costs of rarefying, when there are no systematic differences in library size

In this case, the analysis of unrarefied data using LDM-UR is valid and can be expected to have optimal power for testing hypotheses as it uses all the reads at once. Then, it is of interest to compare the power of LDM-A (the most powerful version of the LDM with rarefaction in our simulations) to LDM-UR. We considered a wide range of the mean library size, which was set to be equal in cases and controls. We considered two levels of rarefaction depth, 25% and 10% of the mean library size (the rarefaction depth was also the minimum library size generated in the data as stated earlier). Note that on average, 90% of the reads are lost in each rarefaction in the second case of rarefaction depth. The results are displayed in Figure We see that rarefaction does lead to loss of power when compared to the analysis of the full data, but the power loss diminishes as the mean library size increases. In the first case of rarefaction depth (25% of mean library size), the power of LDM-A reaches the power of LDM-UR for mean library size above 18K, while in the second case of rarefaction depth (10% of mean library size), the power loss of LDM-A against LDM-UR reduces to less than 3% for mean library size above 18K. Therefore, given the large library sizes from modern sequencing techniques, we expect the loss of power to be small as long as a reasonable rarefaction depth can be used.

### Analysis of the IBD data

We analyzed a subset of the data generated from the RISK cohort, a study of pediatric patients with new-onset Crohn’s Disease (CD) [18] as well as non-IBD controls. For each individual, samples from multiple gastrointestinal locations were collected at the time of diagnosis of CD before treatment initiation, and profiled by 16S rRNA gene sequencing on the Illumina MiSeq platform. The analyses presented here used only data from mucosal tissue biopsies at the rectum site. Further, we filtered out samples with library sizes less than 10,000, which resulted in the loss of 10% samples and allowed for a rarefaction depth of 10,081 reads. In addition, following a common practice, we filtered out taxa that were present in fewer than five samples. Our final data set comprised 267 samples and 2,565 taxa. In our analysis, we defined IBD cases as those with Crohn’s Disease, Indeterminate Colitis, or Ulcerative Colitis; with this choice, we obtained 169 cases and 98 controls. There was an imbalance in the proportion of males by case status (62% in cases, 44% in controls), indicating that sex is a potential confounder. The goal of our analysis was to test for presence-absence association of the rectal microbiome with the IBD status, while controlling for two potential confounders, sex and antibiotic use. It was of interest to test the association at the community level, as well as detecting individual taxa which contributed significantly to the community-level association.

Motivated by other researchers [19] who reported substantial confounding of PERMANOVA results due to library size based on data from the same study, we checked the library size distributions in cases and controls for our selected data. We found that the library size distributions were indeed systematically different (Figure 4). Thus, we rarefied the read count data of all 267 samples to the minimum depth 10,081. We constructed ordination plots (Figure S3) using the Jaccard distance after removing the effects of sex and antibiotic use [9], without rarefaction and with one rarefaction. These plots demonstrated a clear shift in cases compared with controls both before and after rarefaction. Without rarefaction, the two groups seem to separate further apart, corroborating the confounding effect of library size.

**Figure 4.**
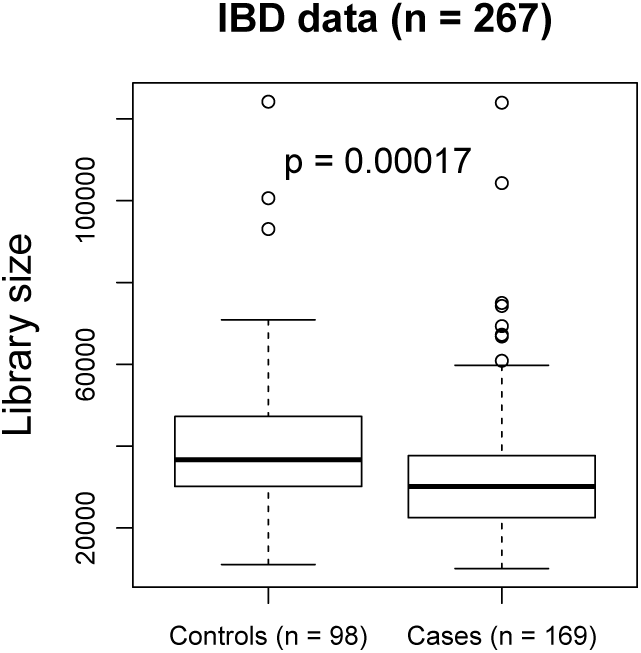
Differential distributions of library sizes in cases and controls of the IBD data.

We applied LDM-A as well as LDM-F(*R*) with the number of rarefactions *R* varying between 1 and 20. The global *p*-values of all tests as shown in Figure 5 (left) are equal to the minimum possible value (i.e., 1*/*5001 where we chose 5,000 to be the maximum number of permutations for the global test), regardless of the number of rarefactions. These *p*-values indicated a very strong presence-absence association of the IBD status with the rectal microbiome at the community level, which is not surprising given the clear shift of case ordinations relative to controls even with only one rarefaction (Figure S3). The number of taxa detected by LDM-F(*R*) (at FDR 10%) increased dramatically as *R* increased from 1 to 5 and stabilized somewhat after 5 (Figure 5, middle), further confirming that at least 5 rarefactions are needed. LDM-A detected the largest number of taxa (469) followed by LDM-F(20) (450). As a comparison, we also applied LDM-UR (without rarefaction), which detected 477 taxa. The Venn diagram in Figure 5 (right) showed that the sets of taxa detected by LDM-F(20) and LDM-A overlap considerably, while the set of taxa by LDM-UR include a large number (94) that do not overlap with any other set, suggesting that these may be false positive findings due to confounding by library size.

**Figure 5.**
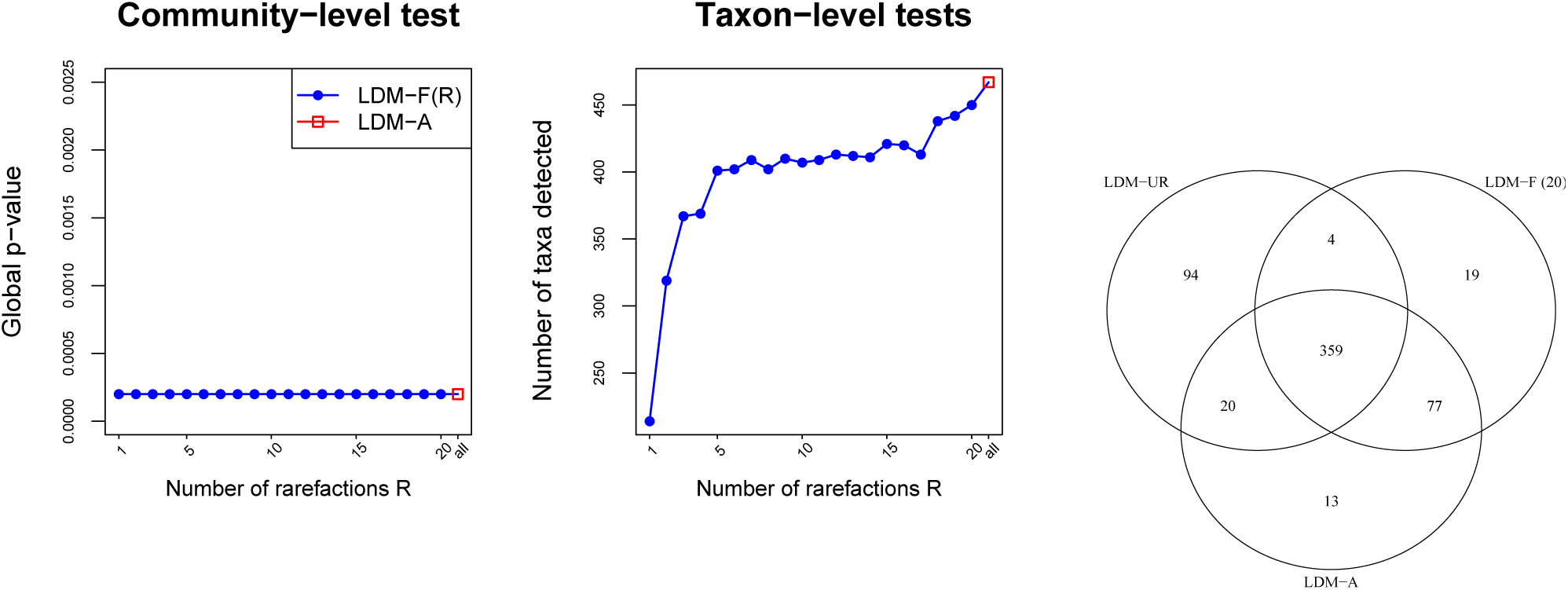
Results in analysis of the IBD data. (Left) Global *p*-value of the community-level test for presence-absence association of the rectal microbiome with the IBD status. (Middle) Number of taxa detected as differentially present between cases and controls at the nominal FDR of 10%. (Right) Venn diagram of taxa detected by LDM-UR, LDM-F(20), and LDM-A.

## Discussion

We have presented two ways of extending the LDM *F*-statistic to aggregate information over multiple rarefactions: the LDM-A, which averages the numerator and denominator RSS terms separately before forming the ratio, and the LDM-F(*R*), which averages the *F*-statistics of *R* rarefaction replicates. We recommend LDM-A over LDM-F(*R*) because it 1) bypasses the selection on the number of rarefaction replicates, 2) yielded more power (and more sensitivity for detecting individual taxa) than LDM-F(*R*) in our simulation studies, and 3) is computationally much more efficient. We have included both methods in our R package `LDM`, which is available on GitHub at https://github.com/yijuanhu/LDM.

Although the use of unrarefied data (LDM-UR) typically gives the best power, when the distributions of library size are similar between cases and controls, the power lost by LDM-A is minimal, as long as the overall sequencing depth is moderately high (*≥*10K) and the rarefaction depth is not too low. Given the robustness of LDM-A to any systematic variation in library size, it seems prudent to use the LDM-A even when there is no obvious difference in the distributions of library size.

The efficiency of the LDM-F(*R*) depends on the rarefaction depth. We note that the LDM-A also depends on the choice of rarefaction depth, both through the parameters *φ*_*ij*_ and in the number of samples retained for the analysis. In general, the guidelines for choosing a rarefaction depth are similar for both methods, as well as other applications that use rarefaction: the depth should be low enough to include most samples, but can exclude samples with anomalously low library sizes.

The LDM is a linear model, which we have applied to binary data. Logistic regression might be seen as a more natural choice for presence-absence data. In fact, the logistic regression model has been adopted for analyzing the zero-part data in several zero-inflated two-part models [20–23]. However, the computational burden of permutation analysis of *R* replicate logistic analyses at each of *J* taxa could be quite extensive, especially when *J* numbers in the thousands. Logistic regression is also problematic when there is separation of the data, which may easily occur at rare taxa, resulting in infinite parameter estimates. Further, for logistic regression, it is not possible in general to analytically average a test statistic over rarefaction replicates for a composite null hypothesis, because any such statistic would depend on parameter estimates which could vary with each replicate. Finally, the computational simplicity of LDM-A and the good performance we observed for presence-absence data argue against using logistic regression.

PERMANOVA is currently the most commonly used method for testing community-level hypotheses about microbiome associations. Like the LDM, PERMANOVA is also based on an *F*-statistic and permutation. It is easy to extend PERMANOVA for testing presence-absence associations by either averaging the numerator and denominator RSS terms that comprise the *F*-statistic separately, or by averaging the *F* statistic directly. However, it seems impossible to calculate a PERMANOVA equivalent of the LDM-A approach in general, because the RSS terms in PERMANOVA are based on a (squared) distance matrix, each entry of which is generally not a linear function of every taxon (with exceptions such as the Euclidean distance matrix). We plan to report our findings on aggregating PERMANOVA over rarefaction replicates in a separate manuscript.

## Conclusions

Analyses of presence-absence associations in the microbiome are generally confounded by the variation of library size. Although rarefaction can be employed to control this bias, it is at the potential cost of information loss as well as the introduction of a stochastic component into the analysis. We presented a novel extension of the LDM for testing presence-absence associations, which is robust to any variation of library size due to the use of rarefaction, statistically efficient due to the aggregation of all rarefied taxa count tables, and computationally efficient due to the analytical (non-stochastic) approach to rarefaction and aggregation. This is currently the only method that provides both the community-level and taxon-level tests of presence-absence associations within one framework.

## Consent for publications

Not applicable.

## Availability of data and materials

The R package `LDM` is available on GitHub at https://github.com/yijuanhu/LDM in formats appropriate for Macintosh, Linux, or Windows systems. The IBD data are available on Qiita at https://qiita.ucsd.edu/study/description/1939.

## Authors’ contributions

YJH conceived the study, developed the method, performed simulation studies, analyzed the data, and wrote the manuscript. GAS conceived the study, developed the method, and wrote the manuscript. AL analyzed the data. All authors read and approved the final manuscript.

## Acknowledgments

Not applicable.

## Ethics approval and consent to participate

This study only involved secondary analyses of existing, de-identified datasets; as such it does not require separate IRB consent.

