## Supplemental Figures for "A Rarefaction-Based Extension of the LDM for Testing Presence-Absence Associations in the Microbiome"

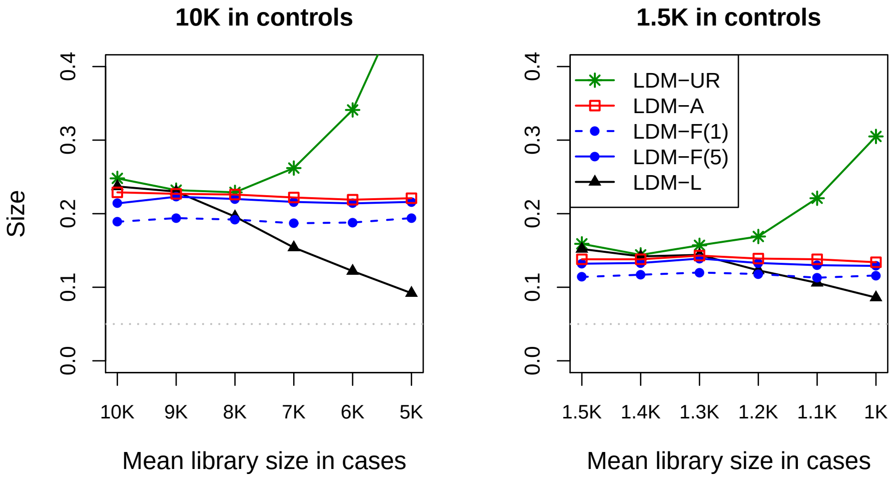


**Figure S1.** Size of the global test, when the confounder $C$ is not controlled for. The gray dotted line represents the nominal significance level of 0.05.

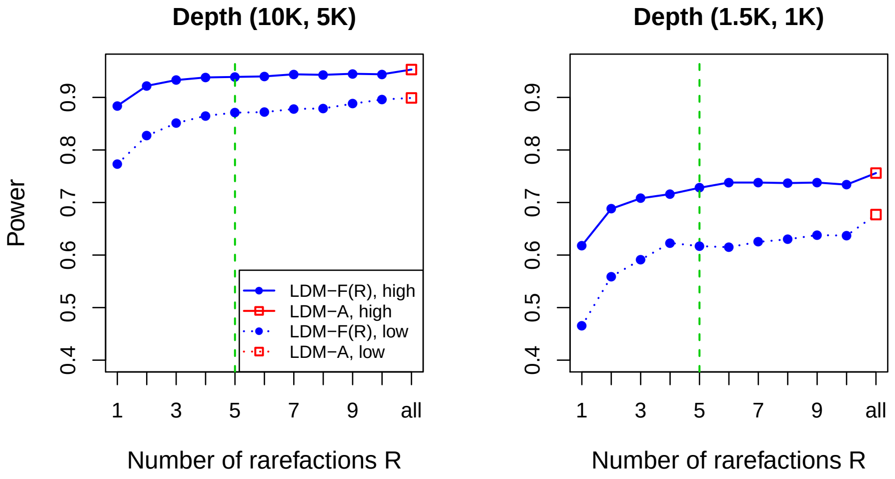


**Figure S2.** Power of the global test by LDM-F($R$) with increasing number of rarefactions and LDM-A with all rarefactions. The solid and dashed lines correspond to settings with high and low rarefaction depths (2.5K vs. 0.5K in the high-throughput setting; 0.5K vs. 0.25K in the low-throughput setting), respectively. The effect size $\beta$ was fixed at 0.5. The green vertical dashed line represents 5 rarefactions.

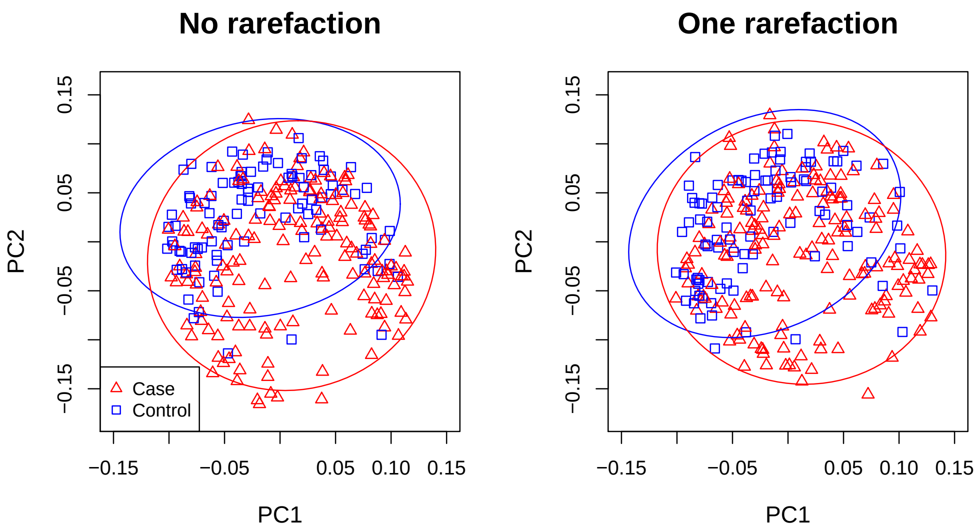


**Figure S3.** Ordination (based on the Jaccard distance after projecting off sex and antibiotic use variables) of the samples in the IBD data, with and without rarefaction. The ellipses are 90% confidence limits for case and control clusters.
